# Coarse-grain simulations on NMR conformational ensembles highlight functional residues in proteins

**DOI:** 10.1101/532507

**Authors:** Sophie Sacquin-Mora

**Affiliations:** Laboratoire de Biochimie Théorique, CNRS UPR9080, Institut de Biologie Physico-Chimique, 13 rue Pierre et Marie Curie, 75005 Paris, France

**Keywords:** NMR conformational ensembles, proteins mechanics, coarse-grain simulations, elastic network model, protein gating residues

## Abstract

Dynamics are a key feature of protein function, and this is especially true of gating residues, which occupy cavity or tunnel lining positions in the protein structure, and will reversibly switch between open and closed conformations in order to control the diffusion of small molecules within a protein’s internal matrix. Earlier work on globins and hydrogenases have shown that these gating residues can be detected using a multiscale scheme combining all atom classic molecular dynamics simulations and coarse grain calculations of the resulting conformational ensemble mechanical properties. Here we show that the structural variations observed in the conformational ensembles produced by NMR spectroscopy experiments are sufficient to induce noticeable mechanical changes in a protein, which in turn can be used to identify residues important for function and forming a *mechanical nucleus* in the protein core. This new approach, which combines experimental data and rapid coarse-grain calculations and no longer needs to resort to time-consuming all-atom simulations, was successfully applied to five different protein families.

## 1. Introduction

Far from being static, rigid objects, proteins are highly flexible molecules, and understanding this flexibility is essential for understanding protein function [1, 2]. Interestingly, protein conformational diversity and dynamics do not always involve large-scale protein motions, and function can arise from small structural changes [3]. In their work investigating the conformational changes between the apo and ligand-bound state in 60 enzymes, Gutteridge and Thornton showed that over 90% of the studied proteins presented root mean square deviations (RMSDs) below 2 Å [4]. These small conformational variations are nonetheless sufficient to support enzymatic activity, and inversely, slightly disrupting a protein’s structure can greatly affect its biological function [5, 6]. As a consequence, current *de novo* protein design strategies do not only involve the production of a particular fold, but also need to take into account protein dynamics, if one wants to achieve a specific function [7–9].

Molecular gates represent a particular class of dynamic structures that play an important role in protein activity. They can be defined as a system, involving individual residues, loops or secondary structure elements, that can reversibly switch between open and closed conformations, and that control the passage of small molecules into and out of the protein structure [10, 11]. Gating residues, lining internal channels connecting a buried active site to the surface, or two distant active sites within a single protein, are a common feature in the protein world, with more than 60% of the enzymes in the Catalytic Site Atlas [12] presenting channels at least 15 Å long [13], and as shown by the 70 examples discussed in a review by Gora et al. [11]. The dynamic properties of these tunnel lining residues play a key role in controlling catalytic efficiency and enzymes substrate specificity [14, 15], and they can be seen as a new category of functional residues, that take a part in regulating biological processes without being directly involved in the catalytic reaction. As a consequence they have attracted growing interest among the drug design community over the recent years [16].

From a theoretical point of view, numerous tools are nowadays available for investigating protein dynamics [14, 17]. Classic all-atom Molecular Dynamics (MD) simulations represent a common strategy, sometimes combined with experimental approaches [18, 19], which can however be computationally costly when working on large systems or modeling events taking place on a long time-scale [20]. Another option is to use coarse-grain models, that combine simplified protein representations and energy functions [21–23]. Elastic Network Models (ENM) [24, 25] in particular have led to many results regarding protein mechanics and dynamics [26–30].

In that perspective, we initially developed the ProPHet (Probing Protein Heterogeneity) program, which models proteins as an elastic network, to investigate protein mechanical properties on the residue level [31]. The rigidity profiles produced by ProPHet then turned out to be a remarkable tool for linking a protein structure to its dynamic properties and its function, showing how small conformational changes within a protein could induce large, non trivial, mechanical effects and in turn impact its activity [32–34]. In earlier sudies on globins [35] and [NiFe]-hydrogenases [36], we also showed that the mechanical variations (calculated with ProPHET) observed in the conformational ensemble produced by classical equilibrium MD simulations can be used to identify a *mechanical nucleus* (MN) in the protein, which is a cluster formed by cavity lining residues occupying conserved positions along the protein sequence. In both cases, MN residues appear to be gating residues, oscillating between rigid and flexible states along the MD trajectory and controlling ligand migration in the vicinity of the active site.

Since its initial developments a few decades ago, NMR spectroscopy has turned out to be a powerful tool for exploring the structure and dynamics of proteins in solution [18, 37–39]. In particular, proteic entries in the Protein Data Bank [40] (PDB) produced by solution NMR often provide a conformational ensemble containing between 10 and 30 structural models instead of a single structure. In this work, we used these conformational ensembles to investigate the mechanical properties in five globular protein families using CG simulations performed by ProPHet. The conformational variations in the experimental NMR ensembles are sufficient to induce large mechanical changes between models from a single PDB entry. Combined with sequence alignements, the coarse-grain calculations based on the NMR structural models permit us to identify MN residues in the proteins without resorting to computationally costly all-atom MD simulations, thus highlighting potential mutational targets in a drug-design perspective.

## 2. Material and Methods

### NMR conformational ensembles

We searched the PDB for proteins where NMR conformational ensembles, comprising a minimum of ten models, are available for at least three different species. This led to the selection of 19 PDB entries belonging to five protein families with various folds listed in table 1: Globins, dihydrofolate reductase (DHFR), cyclophilin A (CPA), peptidyl-tRNA hydrolase (PTH), and parvulin. Each NMR ensemble comprises between 10 and 40 models, and the most representative ones (between 2 and 7 models) were selected using the OLDERADO webserver, http://www.ebi.ac.uk/pdbe/nmr/olderado/ [41]. If the NMR structure was not listed online, we used the *Ensemble cluster* tool, which employs the same method as OLDERADO [42], in the UCSF-Chimera program, https://www.cgl.ucsf.edu/chimera/ [43], to determine the most representative models. Both the ChannelsDB database, https://webchemdev.ncbr.muni.cz/ChannelsDB/index.html [44], and the CASTP webserver, http://sts.bioe.uic.edu/castp/index.html?2r7g [45], were used to identify tunnel and cavity lining residues within all these protein structures.

**Table 1:**
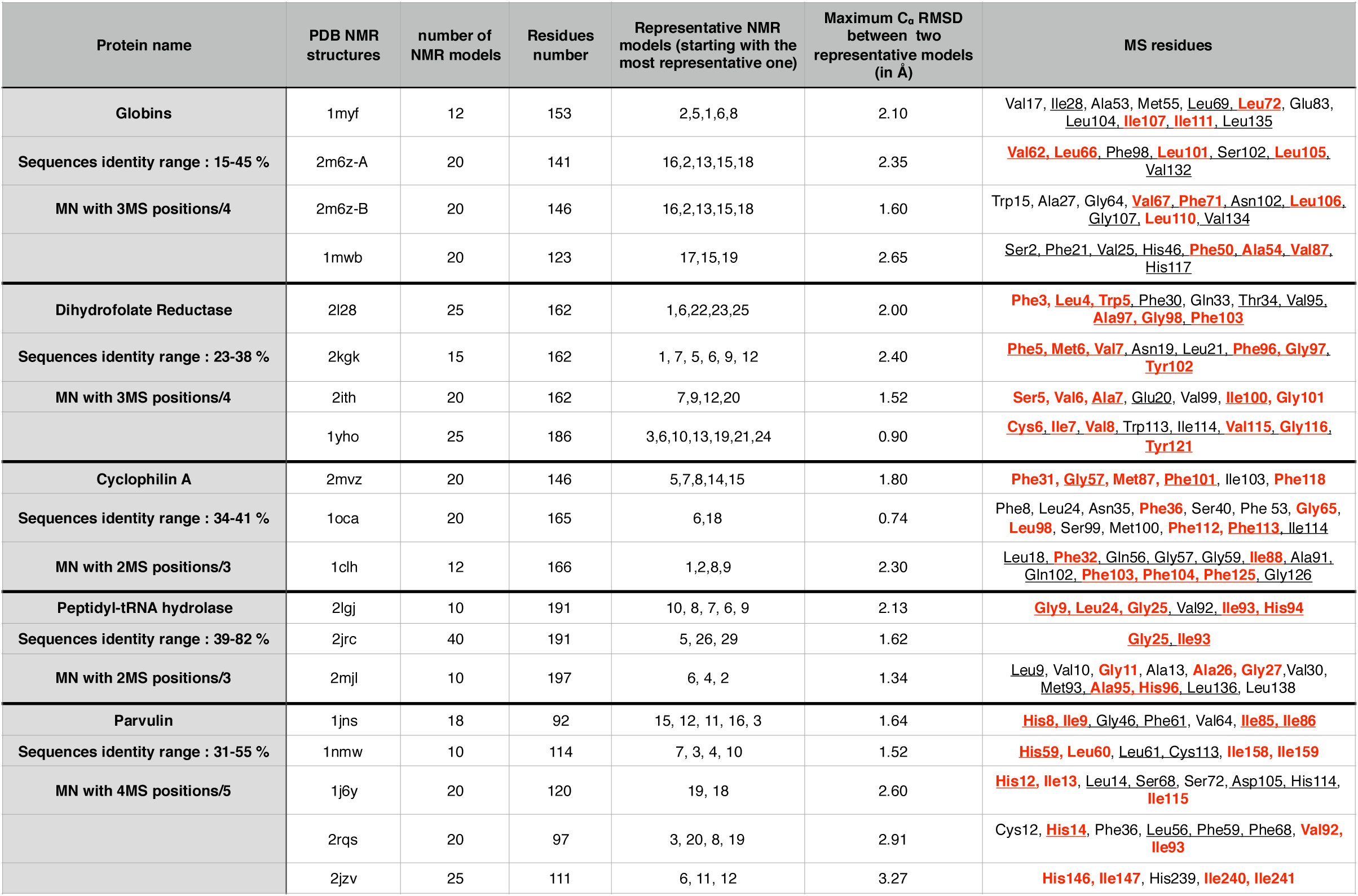
List of the NMR structural ensembles used in this work. In the last column, cavity or tunnel lining residues are underlined and mechanical nucleus residues are highlighted in bold red.

### Coarse-grain Simulation

Coarse-grain Brownian Dynamics (BD) simulations were performed using the ProPHet program, http://bioserv.rpbs.univ-paris-diderot.fr/services/ProPHet/ [33, 34, 46], on all the representative models listed in table 1. Diverging from most common CG models, where each residue is described by a single pseudo-atom [47], ProPHet uses a more detailed model enabling different residues to be distinguished, in which the amino acids are represented by one pseudo-atom located at the C*α* position, and either one or two (for larger residues) pseudo-atoms replacing the side-chain (with the exception of Gly) [48]. Interactions between the pseudo-atoms are treated according to the standard Elastic Network Model (ENM) [47], that is, pseudoatoms closer than the cutoff parameter, *Rc* = 9 Å, are joined by Gaussian springs that all have identical spring constants of *γ* = 0.42 N m^−1^ (0.6 kcal mol^−1^ Å^−2^). The springs are taken to be relaxed for the starting conformation of the protein, i. e. its crystallographic structure. Mechanical properties are obtained from 200,000 BD steps at 300 K. The simulations are analyzed in terms of the fluctuations of the mean distance between each pseudo-atom belonging to a given amino acid and the pseudo-atoms belonging to the remaining protein residues. The inverse of these fluctuations yields an effective force constant *k_i_* that describes the ease of moving a pseudo-atom *i* with respect to the overall protein structure:

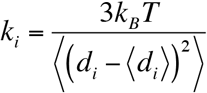

where 〈〉 denotes an average taken over the whole simulation and *d_i_*=〈*d_ij_*〉_*j*\*_ is the average distance from particle *i* to the other particles j in the protein (the sum over *j*\* implies the exclusion of the pseudo-atoms belonging to the same residue as i). The distances between the C*α* pseudo-atom of residue *k* and the C*α* pseudo-atoms of the adjacent residues *k−1* and *k+1* are excluded, since the corresponding distances are virtually constant. The force constant for each residue *k* in the protein is the average of the force constants for all its constituent pseudo-atoms *i*. We will use the term *rigidity profile* to describe the ordered set of force constants for all the residues in a given protein. Note that, following earlier studies which showed how prosthetic groups, such as hemes, or small ligands had little influence on calculated force constants [31, 49], we chose not to include these in the protein representations. This enables us to study the proteins intrinsic mechanical properties independently of the nature and position of any bound ligand.

Following a procedure developed during earlier work on globins [35] and hydrogenases [36], we compared the rigidity profiles obtained for the selected representatives models. The models extracted from the NMR conformational ensembles do not present important structural variations, with Cα rmsds between two models from the same PDB entry being always inferior to 3Å (except for the structure of parvulin domain from *S. aureus*, 2jzv, which presents both a flexible N-terminal tail and a flexible loop). However, these small structural changes are nonetheless sufficient to induce noticeable variations in the mechanical properties of a limited number of residues. We made a systematic pairwise comparison of all the rigidity profiles produced for the representative models from a single PDB entry and, for each residue in the protein, kept the maximum value that could be observed for its force constant variation. We then use the resulting max(|Δk|) profiles to determine *mechanically sensitive (MS) residues*, i. e. residues presenting the largest force constant variations between two structural models. MS residues are defined as residues whose maximum force constant variation satisfies the following criterion:

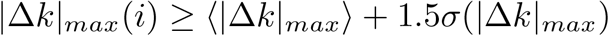

with 〈|Δk|_max_〉 being the average value of the maximum force constant variation over the whole protein sequence and σ(|Δk|_max_) its variance. This procedure leads to the selection of roughly 5-10% of the residues for each one of the NMR structures that are listed in Table 1.

### Sequence alignments

Sequences from pdb entries belonging to the same protein family were aligned using the online MUSCLE tool, https://www.ebi.ac.uk/Tools/msa/muscle/ [50], with the default parameters. Despite the important mechanical similarity that can be observed in the rigidity profiles within a single protein family, the corresponding sequences display relatively low identities ranging from 15% to 82% (see table 1). Mechanically sensitive residues that were identified using the protocol described in the previous section are highlighted in bold and color along the protein sequences in Figure 3a and all throughout the electronic supplementary material.

## 3. Results

### Globins

Four NMR structures were selected in the PDB for the globin family: Myoglobin (Mgb) from *P. catodon* (PDB entry 1myf), truncated hemoglobin (Tr. Hb) from *S. sp. PCC 6803* (1mwb), and the α and β chains from human hemoglobin, Hb (2m6z). The force constant profiles for the main representative model of each protein are shown in Figure 1. Similar to what has been observed in previous studies [35, 46], the analogous aspects of the profiles reflect the common globin fold, with α-helices appearing as grouped rigidity peaks along the protein sequence. The max(Δk) profiles resulting from the systematic pairwise comparison of the force constants calculated for different representative models for a given NMR conformational ensemble are shown in Figure 2 and residues with max(Δk) values above the threshold given in the Material and Methods section (indicated by the red horizontal line) are defined as mechanically sensitive residues, since their mechanical properties are the most impacted by the small conformational variations observed between the NMR models. The MS residue positions are highlighted in red on the multiple sequence alignement shown in Figure 3a and we can see that most of these positions are common to one or more globins. Four positions in particular, that are framed and highlighted in bold green in Figure 3a, E11, E15, G8 and G12 (using the standard numbering for globins that indicates the residue position along helices E and G) correspond to MS residues in three chains out of four. We can see from Figure 3b that these residues form a central cluster right at the heart of the globin fold, with their side chain tightly interacting, and they all correspond to tunnel or pocket lining residues. These four positions also exactly concurs with the mechanical nucleus of globins which was determined in our earlier work using a similar approach, with the alignement of six globin sequences, and the MS residues determination based on clusterized representative structures from trajectories produced by all-atom classical Molecular Dynamic simulations [35]. A search in the literature shows that the mechanical nucleus residues play an important part in protein function by controlling the ligand diffusion process. Positions E11 and G8 were the object of numerous mutational studies in myoglobin [51–53], E11 and E15 are thought to regulate ligand migration in tr. Hb by acting as gate-opening molecular switches [54–57], and in human Hb, gating movements of the leucine residue in G12 governs the hopping of gaseous ligand from and to different binding sites [58]. In neuroglobin, the mechanical properties for all four MN residues have also been shown to be particularly sensitive to pressure changes in high-pressure crystallography [59].

**Figure 1:**
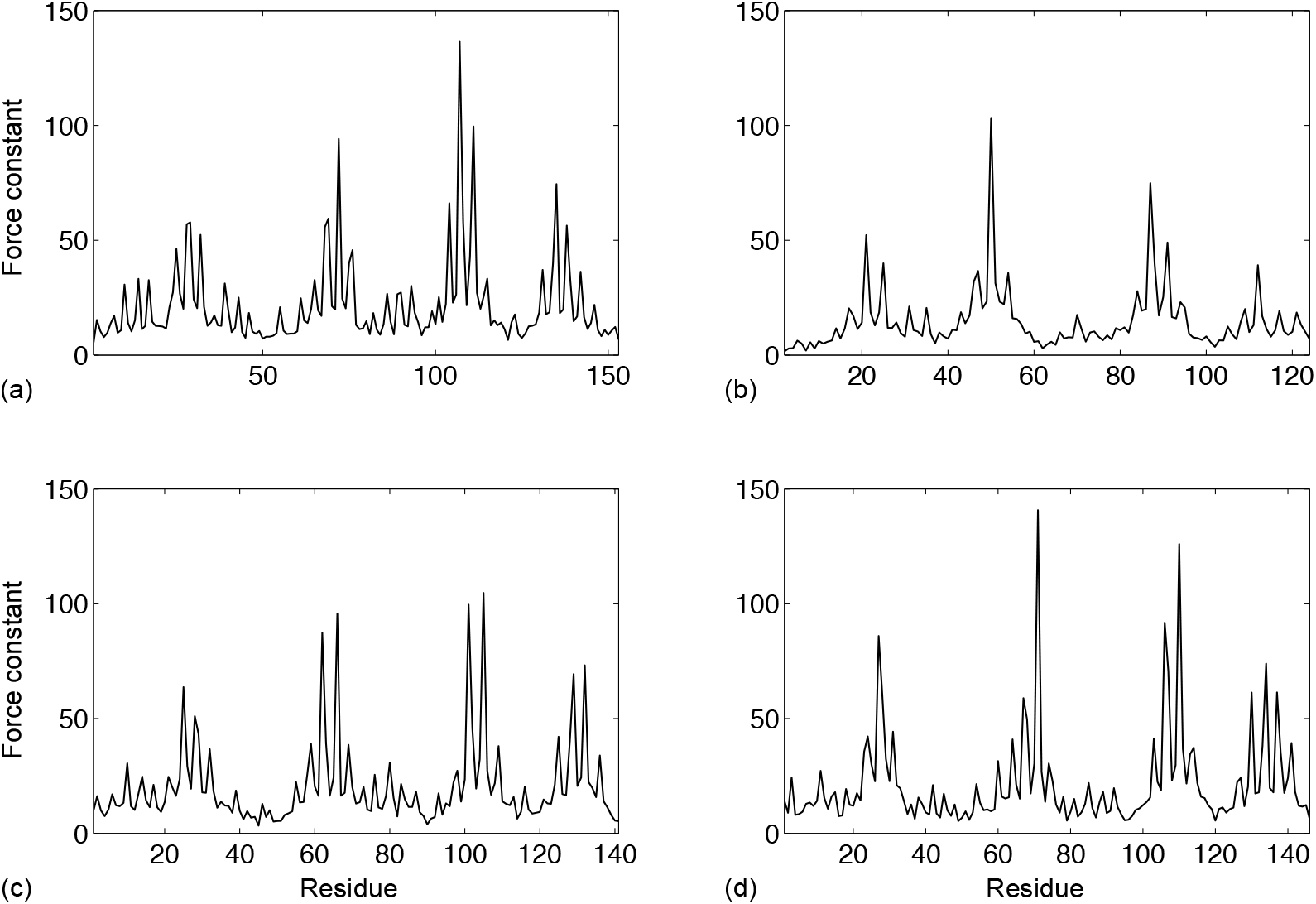
Rigidity profiles (in kcal.mol^−1^.Å^−2^) of the first NMR representative model for the four globin chains under study. (a) Mgb, (b) Tr. Hb, (c) αHb, (d) βHb.

**Figure 2:**
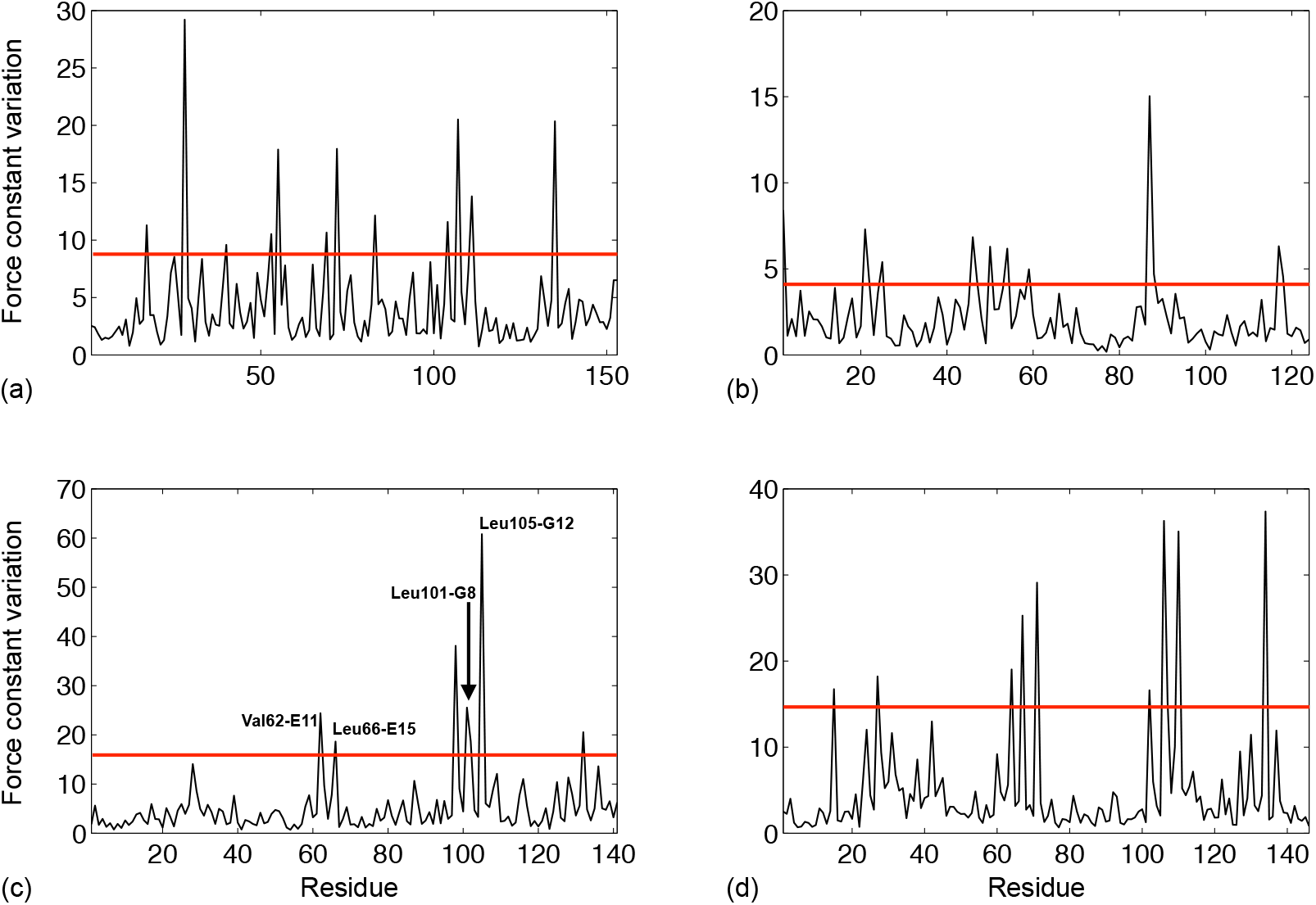
Maximum variation (in kcal.mol^−1^.Å^−2^) of the force constant observed upon pairwise comparison of the NMR representative models. (a) Mgb, (b) Tr. Hb, (c) αHb, (d) βHb. The red horizontal line indicates the threshold value used (following the criterion given in the Material and Methods section) for the selection of mechanically sensitive residues listed in table 1.

**Figure 3:**
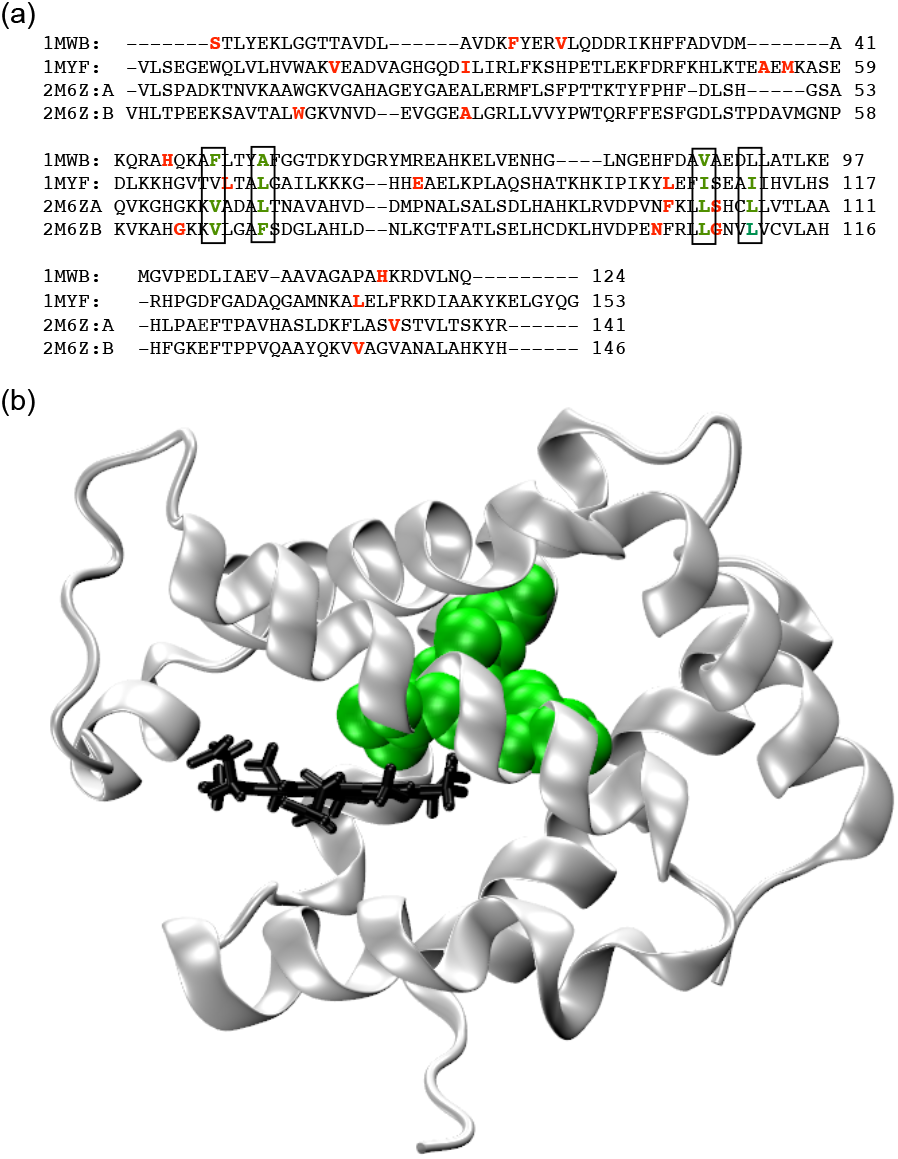
(a) Multiple sequence alignment for the globin structures; mechanically sensitive residues are shown in bold red, and residues belonging to the mechanical nucleus are framed and highlighted in bold green. (b) Cartoon representation of myoglobin with the heme group shown in black and the mechanical nucleus residues as green van der Waals spheres. Figure 3a and Figures 4–7 were prepared using Visual Molecular Dynamics^78^.

### Dihydrofolate reductase

Four NMR conformational ensembles were found in the PDB for dihydrolate reductases (DHFR) from *H. sapiens* (1yho), *L. casei* (2l28), *B. anthracis* (2kgk), and *H. volcanii* (2ith). The combination of coarse-grain mechanical calculations and multiple sequence alignement leads to the identification, when keeping only MS residues that are present in at least 3 out of 4 sequences (see the alignment in Figure SI-1), of a six-residues mechanical nucleus comprising Cys6, Ile7, Val8, Val115, Gly116 and Try121 (using the human sequence numbering) shown in Figure 4. All residues from the mechanical nucleus are listed as tunnel lining or cavity lining in ChannelsDB or CASTP and lie on the interface between the subdomains defined in ref. [60]. A search in the experimental literature shows residues 6-8 belong to the protein initial β-strand, that contains residues critical for ligand and cofactor binding [61, 62], Phe96 from the BaDHFR (corresponding to Val115 in the human DHFR) is also involved in ligand biding [62] and so is Ile94, its equivalent in the *E. coli* DHFR variant [63]. Interestingly, the F98Y mutation in DHFR from *S. aureus* (which corresponds to a mutation of Tyr121 in human DHFR) is responsible for an important loss in TMP affinity, due to Tyr98 forming a hydrogen bond with Leu5 (Ile7 in human), thus preventing it from binding the TMP ligand [64]. The same mutation has also been investigated in DHFR from *M. tuberculosis*, with the Y100F mutant displaying low affinity for classical DHFR inhibitors [65].

**Figure 4:**
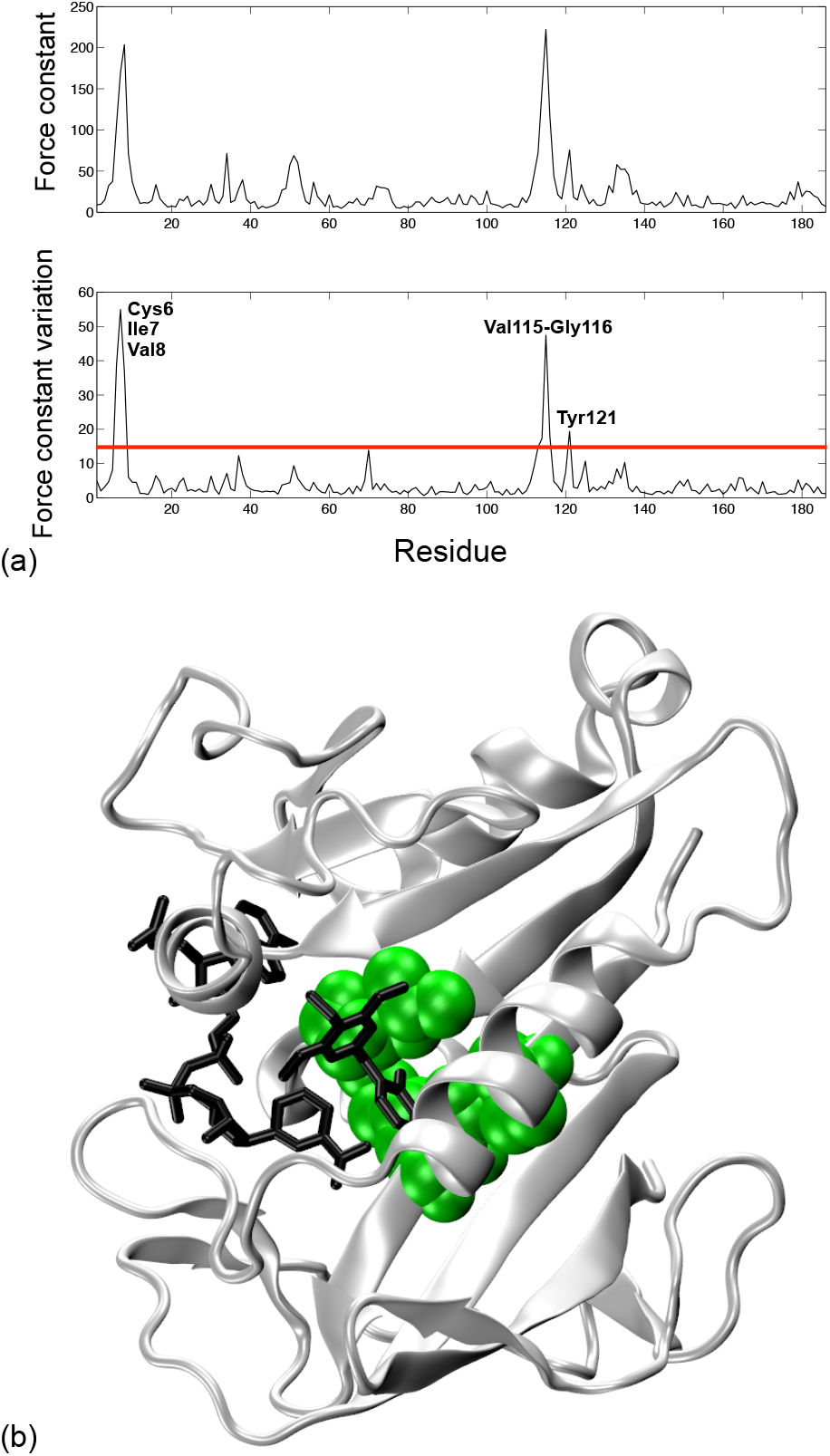
(a) Upper black line, rigidity profile of the first NMR representative model of human DHFR. Lower black line, maximum variation of the force constant observed upon pairwise comparison of the NMR representative models for human DHFR, the

### Cyclophilin A

Coarse-grain calculations were made on three NMR conformational ensembles for cyclophilin A from *H.sapiens* (1oca), *G. kaustophilus* (2mvz) and *E. coli* (1clh). The consensus mechanical nucleus is formed by six residues (Phe36, Gly65, Leu98, Phe112, Phe113 and Phe129, using the human sequence numbering) that correspond to MS positions in at least 2 out of the 3 sequences (see Figure SI-1 for the alignment) and is shown in Figure 5. The ChannelsDB entry for 3rdd, which corresponds to a crystallographic structure for human CPA, indicates that Phe113 is a tunnel lining residues, and CASTP calculations identify all the MS residues in the *E. coli* CPA as cavity lining residues. From the experimental literature, we find that the ligand binding pocket in human CPA comprises residues Gly65, Leu98, Phe112 and Phe113, and Phe36, Phe112 and Phe129 belong to the protein aromatic core [66]. In a study using NMR relaxation experiments [67], Eisenmesser et al. showed how the protein motions play an essential part in its catalytic activity, and this catalytic activity is severely diminished for the F113W mutant. In addition, the large variations in rigidity observed for Leu98, Ser99 and Phe113 in human CPA concur with the fact that these residues display the largest differences in NMR chemical shifts between the enzyme’s major and minor conformers. A mutational study on Ser99 (which lies outside of the catalytic site), designed to stabilized the previously hidden minor conformation, causes a large decrease in the catalytic rate [68]. In the E. coli CPA, residues Ile88, Phe103 and Phe125 belong to the protein hydrophobic core, but are not directly involved in its isomerase activity [69]. Interestingly, MS residues Ala91 and Phe103 are located on both extremities of the B5-B6 surface loop, which overlooks the active site. As a consequence, mechanical variations in these two residues are likely to impact the loop conformational variability, and therefore the ligand’s access to the catalytic site. A similar effect is also observed in the CPA from *G. kaustophilus*, with residues Gly57, Met87 and Phe101 lying on the edge of active site loops [70].

**Figure 5:**
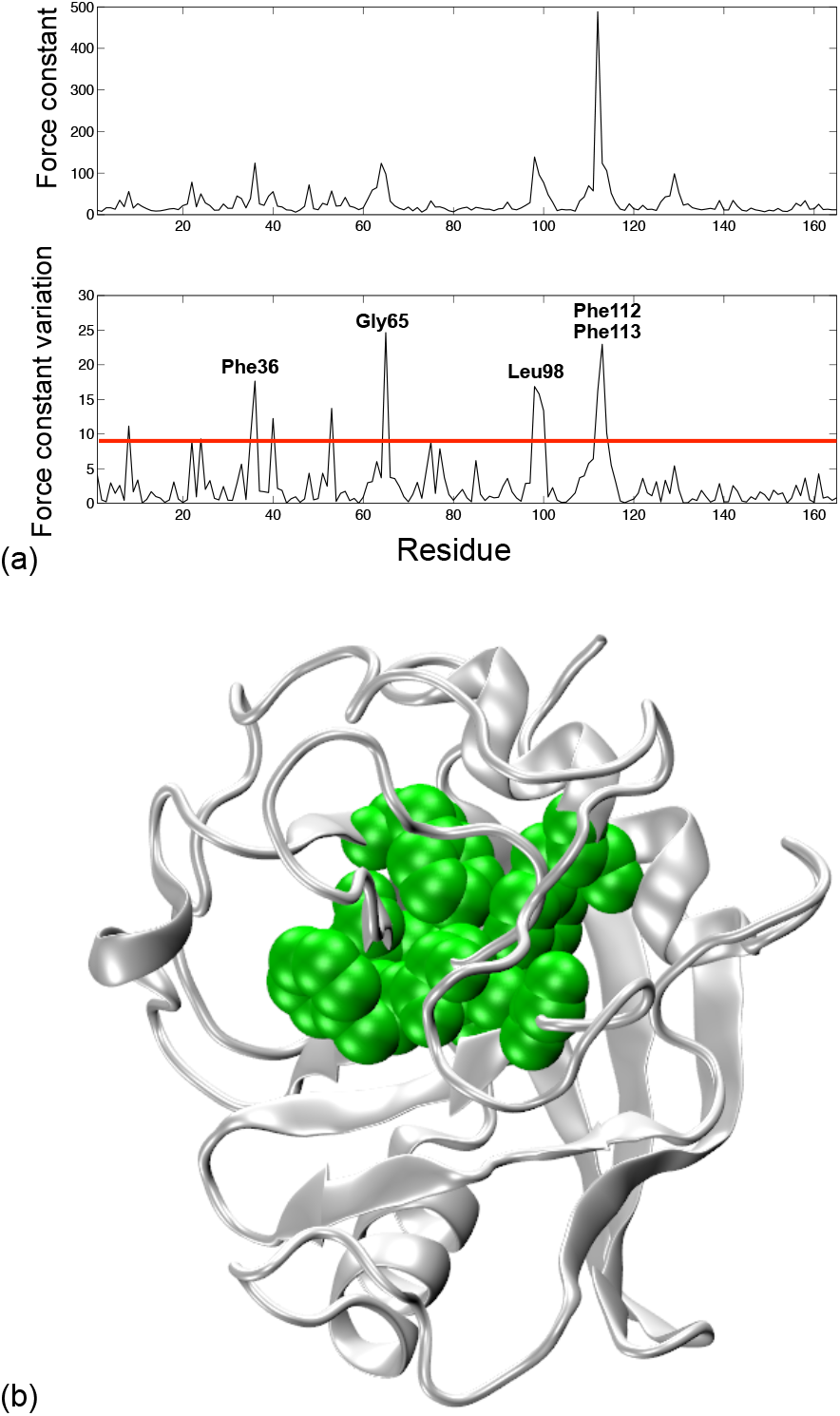
(a) Upper black line, rigidity profile of the first NMR representative model of human CPA. Lower black line, maximum variation of the force constant observed upon pairwise comparison of the NMR representative models for human CPA, the red horizontal line indicates the threshold value used for the selection of mechanically sensitive residues listed in table 1. (b) Cartoon representation of human CPA with the mechanical nucleus residues as green van der Waals spheres.

### Peptidyl-tRNA hydrolase

Three NMR conformational ensembles were found in the PDB for peptidyl-tRNA hydrolase (Pth) from *M. tuberculosis* (2jrc), *M. smegmatis* (2lgj) and *V. cholerae* (2mjl), which lead to the identification of a five-residues mechanical nucleus formed by Gly11, Ala26, Gly27, Ala95 and His96 (using the *V. cholerae* sequence numbering) shown in Figure 6, with MN residue occupying MS positions in at least 2 out of the 3 sequences (see Figure SI-1 for the sequence alignment). All the mechanically sensitive residues listed in table 1 for the 2jrc and the 2lgj structures, are either tunnel lining or cavity lining residues in ChannelsDB or CASTP. The α/β globular fold for Pth comprises an active site channel connecting the surface of the molecule to the central β-sheet core [71]. In the *M. tuberculosis* variant, mechanical nucleus residues are not directly involved in the Pth catalytic activity (unlike the neighboring His24 and Asp97), however, Gly11 and Gly27 are strictly conserved in this proteic family, while Ala95 occupies and hydrophobic positions, and all residues are lining the substrate binding channel [72]. In most Pth variants, Ala95 and His96 lie at the base of the *gate loop*, which controls the entrance to the substrate binding cleft [73–75], and has to move away to permit ligand binding in the enzyme’s active site, which means that a perturbation of their mechanical properties is likely to disrupt the protein catalytic activity. In addition, NMR studies on the *E. coli* Pth show that Ala 26, Gly27 and Ala95 are affected by RNA binding to Pth, despite their central position in the protein core [76], due to the interplay between the tRNA binding region and the peptide binding channel [71]. Ala95 also forms van der Waals contacts with docked substrates in the *V. cholerae* Pth [75].

**Figure 6:**
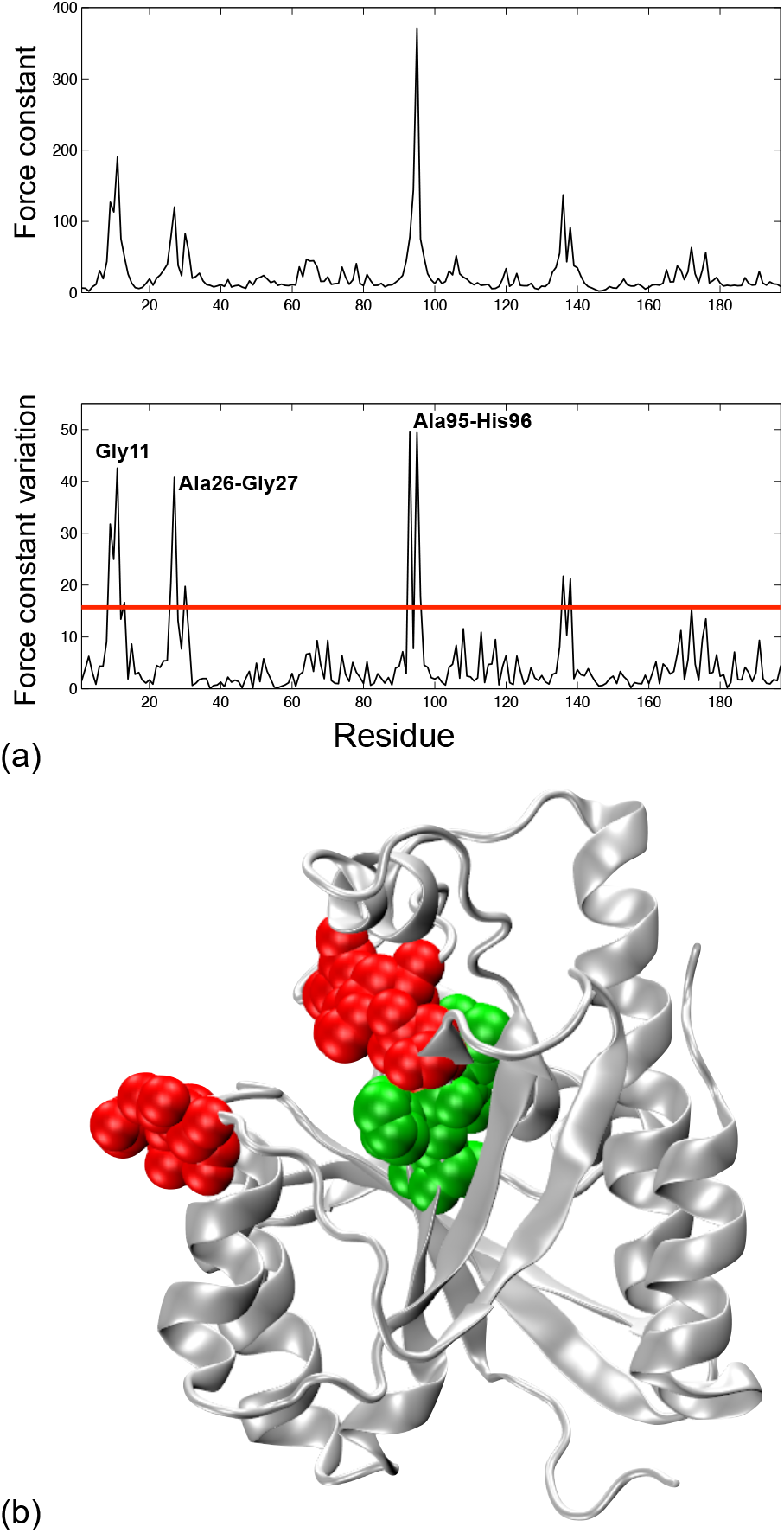
(a) Upper black line, rigidity profile of the first NMR representative model of Pth from *V. cholerae*. Lower black line, maximum variation of the force constant observed upon pairwise comparison of the NMR representative models for Pth from *V. cholerae*, the red horizontal line indicates the threshold value used for the selection of mechanically sensitive residues listed in table 1. (b) Cartoon representation of Pth from *V. cholerae* with the mechanical nucleus residues as green van der Waals spheres and the catalytic residues (at the entrance of the peptide binding cleft) as red van der Waals spheres.

### Parvulin

Five NMR conformational ensembles were found in the PDB for human parvulin (1nmw), and variants from *E. coli* (1jns), *A. thaliana* (1j6y), *C. symbiosum* (2rqs) and *S. aureus* (2jzv), thus leading to the identification of a four-residues mechanical nucleus formed by His59, Ile60, Il158 and Ile159 (using human sequence numbering) shown in Figure 7 (see Figure SI-1 for the sequence alignment). In parvulin from *E. Coli*, all the mechanical nucleus residues are listed as tunnel binding by ChannelsDB and cavity lining by CasTP (see table1). Structural studies on human parvulin highlight the functional importance of the two conserved histidines His59, His157 and Ile159, which are located in the enzyme’s active center [77–79]. Interestingly, later mutational studies on the His59/His157 pair showed that theses residues are not directly involved in the parvulin isomerase activity, but more likely contribute to the enzyme’s stability [79]. His8 in *E. coli* parvulin, and His14 from the *C. symbiosum* parvulin (which both correspond to His59 in human parvulin) are located in the protein hydrophobic core and belong to the putative binding pocket [80, 81], in parvulin from *A. thaliana*, His12 displays no chemical shift after binding of the peptide ligand, thus suggesting that this residue is not directly involved in the catalytic cycle [82]. The NMR conformational ensemble obtained for *S. aureus* parvulin highlights the structural variability of the catalytic histidines (His 146 and His239), and the protein hydrophobic core also comprises residue Ile137, Ile240 and Ile241 [83]. A later study combining Molecular Dynamics Simulation and experimentally available S^2^ order parameter for three parvulins suggests that the conserved histidines play a key role in the modulation of the enzyme’s internal dynamics [84].

**Figure 7:**
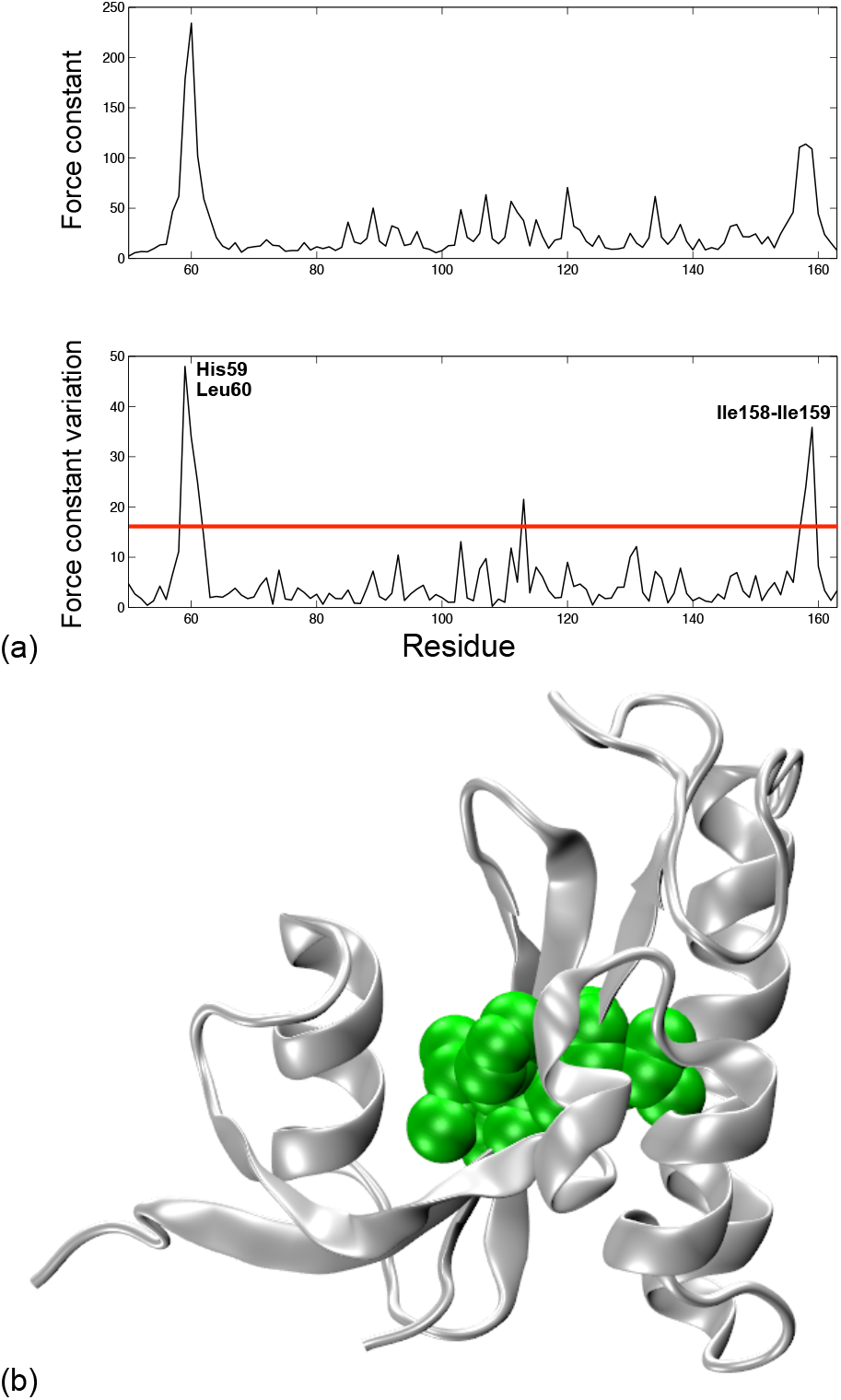
(a) Upper black line, rigidity profile of the first NMR representative model of human parvulin. Lower black line, maximum variation of the force constant observed upon pairwise comparison of the NMR representative models for human parvulin, the red horizontal line indicates the threshold value used for the selection of mechanically sensitive residues listed in table 1. (b) Cartoon representation of human parvulin with the mechanical nucleus residues as green van der Waals spheres.

## 4. Discussion and Conclusion

As shown in earlier work on globins and hydrogenases, protein mechanical properties are a dynamic feature [35, 36]. Due to the robustness of the Elastic Network Model approach [85], a protein’s rigidity profile remains qualitatively the same along time, with its main peaks associated with a given set of residues (that are often likely to be catalytic residues). However, it will also present noticeable variations from one equilibrium structure to the other for another limited number of residues (termed as *mechanically sensitive*, MS, residues), even though the conformational variations between these structures remains negligible. We have shown previously that one could use conformational ensembles produced by classic all-atom Molecular Dynamics trajectories to identify MS residues in a protein, and repeating this approach on several homologous proteins will lead to the definition of a *mechanical nucleus* for this protein family, i. e. a set of conserved positions along the protein sequence that are occupied by MS residues. In the present work, we use protein conformational ensembles obtained by NMR spectroscopy for five distinct protein families, and show that the structural sampling from these ensembles is sufficient to identify a mechanical nucleus comprising between 3 and 6 residues in each group. In the case of globins, this mechanical nucleus concurs perfectly with the globin MN determined using trajectories from MD simulations. Its four constitutive residues have been highlighted as playing a key role in substrate diffusion in the protein matrix in numerous experimental approaches. The MN residues identified in the four other protein families also appear to be excellent candidates for playing a part in protein function as gating residues. Most of them are tunnel or cavity lining residues (underlined residues in Table 1), and for each protein family, experimental evidence, from point mutation studies or evolutionary data, supports the hypothesis that, while not being directly involved in the chemical reaction taking place in the protein active site, the MN residues are nonetheless important for protein function. In particular, by modulating the shape and size of the cavities and tunnels leading to the protein active site, MN residues can impact an enzyme’s substrate selectivity and catalytic efficiency. Therefore, they represent an interesting target for the fine tuning of protein function.

Conformational ensembles built from crystallographic structures can bring a lot of information regarding protein dynamics [86–88]. In order to see wether these could also be used with this approach, we performed mechanical calculations and a pairwise comparison of the rigidity profiles on crystallographic structures of DHFR from three variants, using the PDB IDs listed in [89]: DHFR from *B. anthraci*: chain A from 4alb, 4ale, 4elf, 4elg, 4elh (5 structures), human DHFR, chains A and B from 2w3a, 2w3b and 2w3m (6 structures), DHFR from *E. coli*, chain a from 1rx1, 1rx2, 1rx3, 1rx4, 1rx5, 1rx6, 1rx7, 1rx8, 1rx9 (9 structures). The resulting force constant variation profiles lead to the same mechanical nucleus as when using NMR ensembles, except for Ty121 (using the human sequence numbering), which is no longer a mechanically sensitive residues for any of the three species (see Figure SI-2). This loss in the mechanical nucleus definition is not negligible, since this position has been shown to be functionally important for DHFR from *S. aureus* and *M. tuberculosis* [64, 65].

Another issue with ensembles of crystallographic structures is the choice of representative structures which will be used for mechanical calculations (ideally one should retain around 4-6 models to limit the computational cost of the approach). In the dataset listed in Ref. [89], at least 45 structures are retained for each protein. However, selecting structures that are to mechanically too close can lead to a poor discrimination of mechanically sensitive residues, resulting in a poorly defined mechanical nucleus. As an example, we performed mechanical calculations on 10 crystallographic structures from human cyclophilin A (PDB 5noq 5nor 5nos 5not 5nou 5nov 5now 5nox 5noy 5noz). The pairwise comparison of all ten rigidity profiles leads to the same selection of mechanically sensitive residues as when using the NMR conformational ensemble (see the lower black line in Figure SI-3a). However, if one selects structures that are too close from a mechanical point of view (namely 5noq, 5nor, 5nou and 5nov), the residues force constant variations are always below 10 kcal mol^−1^ Å^−2^, thus making the selection of mechanically sensitive residues more difficult (see the red line in Figure SI-3a), a situation that was never observed for any of the 19 NMR ensembles used in this work. Moreover, in the case of structures with low rmsds (below 2.5 Å), the rmsd and the maximum force constant variation between two structures are not correlated, as can be seen on Figure SI-3b and Figure SI-4b. Which means that selecting structures on the basis of their rmsd is no guarantee that they will be sufficiently different from a mechanical point of view.

Altogether, it seems that our methode can also use crystallographic structures. But this will request more work from a potential user, since one has to build an ensemble from several PDB structures, instead of using a single NMR ensemble, and be careful that the mechanical properties of the selected structures vary sufficiently in order to clearly define the mechanically sensitive residues.

Finally, our use of NMR structural ensembles and coarse-grain calculations provides us with a simple and efficient tool for identifying key spots in the protein structure, where point mutations could help modulate the protein enzymatic activity or specificity. While the current study has been limited to small proteins, comprising less than 200 residues, recent progress in NMR spectroscopy on large systems [39] should ensure that experimental conformational ensembles for larger proteins will be accessible in the near future.

## Acknowledgments

This work was supported by the “Initiative d’Excellence” program from the French State (Grant “DYNAMO”, ANR-11-LABX-0011-01).

## Supporting Material

The multiple sequence alignments for the 5 protein families under study are available as supporting information.

---

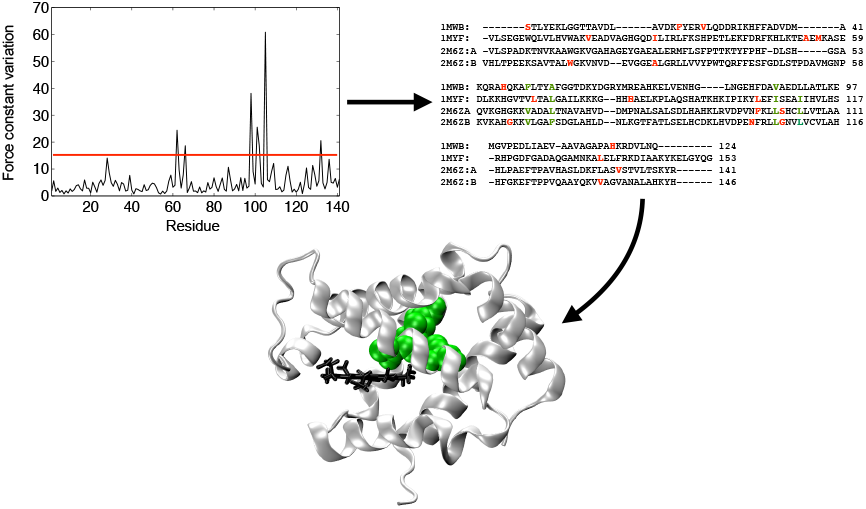
TOC image

