## Supplementary Information for "Coarse-grain simulations on NMR conformational ensembles highlight functional residues in proteins"

Sophie Sacquin-Mora

*Laboratoire de Biochimie Théorique, CNRS UPR9080, Institut de Biologie Physico-Chimique, 13 rue Pierre et Marie Curie, 75005 Paris, France*

**Fig. SI-1:** MUSCLE multiple sequence alignments for the 5 protein families listed in Table 1. Rigid residues are shown in bold red, consensus nucleus residues are in bold green.

CLUSTAL multiple sequence alignment by MUSCLE (3.8)

### Globins

```
1MWB:   -------STLYEKLGGTTAVDL------AVDKFYERVLQDDRIKHFFADVDM-------A 41
1MYF:   -VLSEGEWQLVLHVWAKVEADVAGHGQDILIRLFKSHPETLEKFDRFKHLKTEAEMKASE 59
2M6Z:A -VLSPADKTNVKAAWGKVGAHAGEYGAEALERMFLSFPTTKTYFPHF-DLSH-----GSA 53
2M6Z:B VHLTPEEKSAVTALWGKVNVD--EVGGEALGRLLVVYPWTQRFFESFGDLSTPDAVMGNP 58

1MWB: KQRAHQKAFLTYAFGGTDKYDGRYMREAHKELVENHG----LNGEHFDAVAEDLLATLKE 97
1MYF: DLKKHGVTVLTALGAILKKKG--HHEAELKPLAQSHATKHKIPIKYLEFISEAIIHVLHS 117
2M6ZA QVKGHGKKVADALTNAVAHVD--DMPNALSALSDLHAHKLRVDPVNFKLLSHCLLVTLAA 111
2M6ZB KVKAHGKKVLGAFSDGLAHLD--NLKGTFATLSELHCDKLHVDPENFRLLGNVLVCVLAH 116

1MWB:   MGVPEDLIAEV-AAVAGAPAHKRDVLNQ--------- 124
1MYF:   -RHPGDFGADAQGAMNKALELFRKDIAAKYKELGYQG 153
2M6Z:A  -HLPAEFTPAVHASLDKFLASVSTVLTSKYR------ 141
2M6Z:B  -HFGKEFTPPVQAAYQKVVAGVANALAHKYH------ 146
```

### Dihydrofolate Reductase

```
2ITH: -MELVSVAALAENRVIGRDGELPWPSIPADKKQYRSRVA-------DDPVVLGRTTFESM 52
1YHO: VGSLNCIVAVSQNMGIGKNGDLPWPPLRNEFRYFQRMTTTSSVEGKQNLVIMGKKTWFSI 60
2L28: ---TAFLWAQDRDGLIGKDGHLPW-HLPDDLHYFRAQTV-------GKIMVVGRRTYESF 42
2KGK: -MIVSFMVAMDENRVIGKDNNLPW-RLPSELQYVKKTTM-------GHPLIMGRKNYEAI 51

2ITH: ---RDDLPGSAQIVMSRSERSFSVDTAH-RAASVEEAVDIAASLD----AETAYVIGGAA 104
1YHO: PEKNRPLKGRINLVLSRELK-EPPQGAHFLSRSLDDALKLTEQPELANKVDMVWIVGGSS 119
2L28: P--KRPLPERTNVVLTHQED-YQAQGAV-VVHDVAAVFAYAKQHP----DQELVIAGGAQ 101
2KGK: ---GRPLPGRRNIIVTRNEG-YHVEGCE-VAHSVEEVFELCKN------EEEIFIFGGAQ 100

2ITH: IY--ALFQPHLDRMVLSRVPGEYEGDTYYPEWDAAEWE--------LDAETDHEG--FTL 152
1YHO: VYKEAMNHPGHLKLFVTRIMQDFESDTFFPEIDLEKYKLLPEYPGVLSDVQEEKGIKYKF 179
2L28: IF--TAFKDDVDTLLVTRLAGSFEGDTKMIPLNWDDFTKVSSRT--VEDTNPALT--HTY 155
2KGK: IY--DLFLPYVDKLYITKIHHAFEGDTFFPEMDMTNWKEVFVEKGLTDEKNPYT---YYY 155

2ITH: QEWVRSASSR------- 162
1YHO: EVYEKND---------- 186
2L28: EVWQKKA---------- 162
2KGK: HVYEKQQLEHHHHHHHH 186
```

### Cyclophilin A

```
1CLH: -AKGDPHVLLTTSA---GNIELELDKQKAPVSVQNFVDYV--NSGF-YNNTTFHRVIPGF 53
2MVZ: GSMAKKGYILMENG---GKIEFELFPNEAPVTVANFEKLA--NEGF-YNGLTFHRVIPGF 52
1OCA: MVNPTVFFDIAVDGEPLGRVSFELFADKVPKTAENFRALSTGEKGFGYKGSCFHRIIPGF 60

1CLH: MIQGGGFTEQMQQ-KKPNPPIKNEADNG-LRNT-RGTIAMARTADKDSATSQFFINVADN 110
2MVZ: VSQGGC-PRGNGT-GDAGYTIPCETDNNPHRHV-TGAMSMAH-RGRDTGSCQFFIVHEPQ 108
1OCA: MCQGGDFTRHNGTGGKSIYGEKFEDENFILKHTGPGILSMAN-AGPNTNGSQFFICTAKT 119

1CLH: AFLDHGQRDFGYAVFGKVVKGMDVADKISQVPTHDVGPYQNVPSKPVVILSAKVLP 166
2MVZ: PHLDG-----VHTVFGQVTSGMDVVRTM----------KNGDVMKEVKVFDE---P 146
1OCA: EWLDG-----KHVVFGKVKEGMNIVEAMER-----FGSRNGKTSKKITIADCGQLE 165
```

### Peptidyl t-RNA hydrolase

```
2MJL: MVSQPIKLLVGLANPGPEYAKTRHNAGAWVVEELARIHNVTLKNEPKFFG--LTGRLLIN 58
2LGJ: -MAEPL-LVVGLGNPGPTYAKTRHNLGFMVADVLAGRIGSAFKVHKKSGAEVVTGRL--A 56
2JRC: -MAEPL-LVVGLGNPGANYARTRHNLGFVVADLLAARLGAKFKAHKRSGAEVATGRS--A 56

2MJL: SQELRVLIPTTFMNLSGKAIAALANFYQIKPEEIMVAHDELDLPPGVAKFKQGGGHGGHN 118
2LGJ: GTSVVLAKPRCYMNESGRQVGPLAKFYSVPPQQIVVIHDELDIDFGRIRLKLGGGEGGHN 116
2JRC: GRSLVLAKPRCYMNESGRQIGPLAKFYSVAPANIIVIHDDLDLEFGRIRLKIGGGEGGHN 116

2MJL: GLKDTISKLGNNKEFYRLRLGIGHPGHKDKVAGYVLGKAPAKEQECLDAAVDESVRCLEI 178
2LGJ: GLRSVASALG-TKNFHRVRIGVGRPPGRKDPAAFVLENFTAAERAEVPTIVEQAADATEL 175
2JRC: GLRSVVAALG-TKDFQRVRIGIGRPPGRKDPAAFVLENFTPAERAEVPTICEQAADATEL 175

2MJL:  LMKDGLTKAQNRLHTFKAE---------- 197
2LGJ:  LIAQGLEPAQNTVHAW------------- 191
2JRC:  LIEQGMEPAQNRVHAWKLAAALEHHHHHH 191
```

### Parvulin

```
1JNS: ------------------------AKTAAALHILVK------------------------ 12
2JZV: -----------------GPLGS--DSKKAS-HILIKVKSKKSDKEGLDDK---------- 164
2RQS: -----------------GPMGSMADKIKCS-HILVK------------------------ 18
1NMW: ---------------------GEPARVRCS-HLLVKHSQSRRPSSWRQEK----ITRTKE 83
1J6Y: MGSSHHHHHHSSGLVPRGSHMASRDQVKAS-HILIKHQGSRRKASWKDPEGKIILTTTRE 40

1JNS: --EEKLALDLLEQIKNGADFGKLAKKHS-ICPSGKRGGDLGEFRQGQMVPAFDKVVFSCP 69
2JZV: EAKQKAEEIQKEVSKDPSKFGEIAKKESMDTGSAKKDGELGYVLKGQTDKDFEKALFKLK 224
2RQS: --KQGEALAVQERLKAGEKFGKLAKELSIDGGSAKRDGSLGYFGRGKMVKPFEDAAFRLQ 76
1NMW: EALELINGYIQKIKSGEEDFESLASQFS-DCSSAKARGDLGAFSRGQMQKPFEDASFALR 142
1J6Y: AAVEQLKSIREDIVSGKANFEEVATRVS-DCSSAKRGGDLGSFGRGQMQKPFEEATYALK 99

1JNS: VLEPTGPLHTQFGYHIIKVLYRN 92
2JZV: DGEVSEVVKSSFGYHIIKADK-- 245
2RQS: VGEVSEPVKSEFGYHVIKRLG-- 97
1NMW: TGEMSGPVFTDSGIHIILRTE-- 163
1J6Y: VGDISDIVDTDSGVHIIKRTA-- 120
```

### References were the predicted mechanical nucleus residues are highlighted as functionally relevant residues:

#### Globins: Val62, Leu66, Leu101, Leu105 (for myoglobin, 1myf)

- Quillin, M.L., Li, T.S., Olson, J.S., Phillips, G.N., Dou, Y., Ikedasaito, M., Regan, R., Carlson, M., Gibson, Q.H., Li, H.Y., et al. 1995 Structural and functional-effects of apolar mutations of the distal valine in myoglobin. *J. Mol. Biol.* **245**, 416-436.

- Ishikawa, H., Uchida, T., Takahashi, S., Ishimori, K. & Morishima, I. 2001 Ligand migration in human myoglobin: Steric effects of isoleucine 107(G8) on O-2 and CO binding. *Biophys. J.* **80**, 1507-1517.

- Dantsker, D., Roche, C., Samuni, U., Blouin, G., Olson, J.S. & Friedman, J.M. 2005 The position 68(E11) side chain in myoglobin regulates ligand capture, bond formation with heme iron, and internal movement into the xenon cavities. *J. Biol. Chem.* **280**, 38740-38755.

- Bidon-Chanal, A., Marti, M.A., Crespo, A., Milani, M., Orozco, M., Bolognesi, M., Luque, F.J. & Estrin, D.A. 2006 Ligand-induced dynamical regulation of NO conversion in Mycobacterium tuberculosis truncated hemoglobin-N. *Proteins* **64**, 457-464.

- Bidon-Chanal, A., Marti, M.A., Estrin, D.A. & Luque, F.J. 2007 Dynamical regulation of ligand migration by a gate-opening molecular switch in truncated hemoglobin-N from Mycobacterium tuberculosis. *J. Am. Chem. Soc.* **129**, 6782-6788.

- Lama, A., Pawaria, S., Bidon-Chanal, A., Anand, A., Gelpi, J.L., Arya, S., Marti, M., Estrin, D.A., Luque, F.J. & Dikshit, K.L. 2009 Role of Pre-A Motif in Nitric Oxide Scavenging by Truncated Hemoglobin, HbN, of Mycobacterium tuberculosis. *J. Biol. Chem.* **284**, 14457-14468.

- Mishra, S. & Meuwly, M. 2009 Nitric Oxide Dynamics in Truncated Hemoglobin: Docking Sites, Migration Pathways, and Vibrational Spectroscopy from Molecular Dynamics Simulations. *Biophys. J.* **96**, 2105-2118.

- Savino, C., Miele, A.E., Draghi, F., Johnson, K.A., Sciara, G., Brunori, M. & Vallone, B. 2009 Pattern of Cavities in Globins: The Case of Human Hemoglobin. *Biopolymers* **91**, 1097-1107.

- Colloc'h, N., Sacquin-Mora, S., Avella, G., Dhaussy, A.C., Prange, T., Vallone, B. & Girard, E. 2017 Determinants of neuroglobin plasticity highlighted by joint coarse-grained simulations and high pressure crystallography. *Scientific reports* **7**, 1858.

#### Dihydrofolate reductase: Cys6, Ile7, Val8, Val115, Gly116, Tyr121 (human DHFR, 1yho)

- Kovalevskaya, N.V., Smurnyy, Y.D., Polshakov, V.I., Birdsall, B., Bradbury, A.F., Frenkiel, T. & Feeney, J. 2005 Solution structure of human dihydrofolate reductase in its complex with trimethoprim and NADPH. *J Biomol NMR* **33**, 69-72.

- Beierlein, J.M., Deshmukh, L., Frey, K.M., Vinogradova, O. & Anderson, A.C. 2009 The solution structure of Bacillus anthracis dihydrofolate reductase yields insight into the analysis of structure-activity relationships for novel inhibitors. *Biochemistry* **48**, 4100-4108.

- Bhabha, G., Lee, J., Ekiert, D.C., Gam, J., Wilson, I.A., Dyson, H.J., Benkovic, S.J. & Wright, P.E. 2011 A dynamic knockout reveals that conformational fluctuations influence the chemical step of enzyme catalysis. *Science* **332**, 234-238. (doi:10.1126/science.1198542).

- Frey, K.M., Liu, J., Lombardo, M.N., Bolstad, D.B., Wright, D.L. & Anderson, A.C. 2009 Crystal structures of wild-type and mutant methicillin-resistant Staphylococcus aureus dihydrofolate reductase reveal an alternate conformation of NADPH that may be linked to trimethoprim resistance. *J. Mol. Biol.* **387**, 1298-1308.

- Dias, M.V., Tyrakis, P., Domingues, R.R., Paes Leme, A.F. & Blundell, T.L. 2014 Mycobacterium tuberculosis dihydrofolate reductase reveals two conformational states and a possible low affinity mechanism to antifolate drugs. *Structure* **22**, 94-103. (doi:10.1016/j.str.2013.09.022).

### Cyclophilin A: Phe36, Gly65, Leu98, Phe112, Phe113, Phe129 (human Cyp A, 1oca)

- Ottiger, M., Zerbe, O., Guntert, P. & Wuthrich, K. 1997 The NMR solution conformation of unligated human cyclophilin A. *J. Mol. Biol.* **272**, 64-81.

- Eisenmesser, E.Z., Millet, O., Labeikovsky, W., Korzhnev, D.M., Wolf-Watz, M., Bosco, D.A., Skalicky, J.J., Kay, L.E. & Kern, D. 2005 Intrinsic dynamics of an enzyme underlies catalysis. *Nature* **438**, 117-121.

- Fraser, J.S., Clarkson, M.W., Degnan, S.C., Erion, R., Kern, D. & Alber, T. 2009 Hidden alternative structures of proline isomerase essential for catalysis. *Nature* **462**, 669-673.

- Clubb, R.T., Ferguson, S.B., Walsh, C.T. & Wagner, G. 1994 Three-dimensional solution structure of Escherichia coli periplasmic cyclophilin. *Biochemistry* **33**, 2761-2772.

- Holliday, M.J., Camilloni, C., Armstrong, G.S., Isern, N.G., Zhang, F., Vendruscolo, M. & Eisenmesser, E.Z. 2015 Structure and Dynamics of GeoCyp: A Thermophilic Cyclophilin with a Novel Substrate Binding Mechanism That Functions Efficiently at Low Temperatures. *Biochemistry* **54**, 3207-3217.

### Peptidyl-tRNA hydrolase: Gly11, Ala26, Gly27, Ala95, His96 (Pth from *V. cholerae*, 2mjl)

- Pulavarti, S.V., Jain, A., Pathak, P.P., Mahmood, A. & Arora, A. 2008 Solution structure and dynamics of peptidyl-tRNA hydrolase from Mycobacterium tuberculosis H37Rv. *J. Mol. Biol.* **378**, 165-177.

- Kabra, A., Fatma, F., Shahid, S., Pathak, P.P., Yadav, R., Pulavarti, S.V., Tripathi, S., Jain, A. & Arora, A. 2016 Structural characterization of peptidyl-tRNA hydrolase from Mycobacterium smegmatis by NMR spectroscopy. *Biochim. Biophys. Acta* **1864**, 1304-1314.

- Singh, A., Gautam, L., Sinha, M., Bhushan, A., Kaur, P., Sharma, S. & Singh, T.P. 2014 Crystal structure of peptidyl-tRNA hydrolase from a Gram-positive bacterium, Streptococcus pyogenes at 2.19 A resolution shows the closed structure of the substrate-binding cleft. *FEBS Open Bio* **4**, 915-922.

- Zhang, F., Song, Y., Niu, L., Teng, M. & Li, X. 2015 Crystal structure of Staphylococcus aureus peptidyl-tRNA hydrolase at a 2.25 A resolution. *Acta Biochim Biophys Sin (Shanghai)* **47**, 1005-1010.

- Kabra, A., Shahid, S., Pal, R.K., Yadav, R., Pulavarti, S.V., Jain, A., Tripathi, S. & Arora, A. 2017 Unraveling the stereochemical and dynamic aspects of the catalytic site of bacterial peptidyl-tRNA hydrolase. RNA 23 , 202-216. (doi:10.1261/rna.057620.116).

- Giorgi, L., Bontems, F., Fromant, M., Aubard, C., Blanquet, S. & Plateau, P. 2011 RNA-binding site of Escherichia coli peptidyl-tRNA hydrolase. *The Journal of biological chemistry* **286**, 39585-39594.

### Parvulin: His59, Leu60, Ile158, Ile159 (human parvulin, 1nmw)

- Ranganathan, R., Lu, K.P., Hunter, T. & Noel, J.P. 1997 Structural and functional analysis of the mitotic rotamase Pin1 suggests substrate recognition is phosphorylation dependent. *Cell* **89**, 875-886.

- Bayer, E., Goettsch, S., Mueller, J.W., Griewel, B., Guiberman, E., Mayr, L.M. & Bayer, P. 2003 Structural analysis of the mitotic regulator hPin1 in solution: insights into domain architecture and substrate binding. *The Journal of biological chemistry* **278**, 26183-26193. (doi:10.1074/jbc.M300721200).

- Bailey, M.L., Shilton, B.H., Brandl, C.J. & Litchfield, D.W. 2008 The dual histidine motif in the active site of Pin1 has a structural rather than catalytic role. *Biochemistry* **47**, 11481-11489.

- Kuhlewein, A., Voll, G., Hernandez Alvarez, B., Kessler, H., Fischer, G., Rahfeld, J.U. & Gemmecker, G. 2004 Solution structure of Escherichia coli Par10: The prototypic member of the Parvulin family of peptidyl-prolyl cis/trans isomerases. *Protein science : a publication of the Protein Society* **13**, 2378-2387.

- Jaremko, L., Jaremko, M., Elfaki, I., Mueller, J.W., Ejchart, A., Bayer, P. & Zhukov, I. 2011 Structure and dynamics of the first archaeal parvulin reveal a new functionally important loop in parvulin-type prolyl isomerases. *The Journal of biological chemistry* **286**, 6554-6565.

- Landrieu, I., Wieruszeski, J.M., Wintjens, R., Inze, D. & Lippens, G. 2002 Solution structure of the single-domain prolyl cis/trans isomerase PIN1At from Arabidopsis thaliana. *J. Mol. Biol.* **320**, 321-332.

- Heikkinen, O., Seppala, R., Tossavainen, H., Heikkinen, S., Koskela, H., Permi, P. & Kilpelainen, I. 2009 Solution structure of the parvulin-type PPIase domain of Staphylococcus aureus PrsA--implications for the catalytic mechanism of parvulins. *BMC Struct Biol* **9**, 17.

- Czajlik, A., Kovacs, B., Permi, P. & Gaspari, Z. 2017 Fine-tuning the extent and dynamics of binding cleft opening as a potential general regulatory mechanism in parvulin-type peptidyl prolyl isomerases. *Scientific reports* **7**, 44504.

**Figure SI-2:** (a) Upper black line, rigidity profile for the crystallographic structure of human DHFR (pdb 2w3a, chain A). Lower black line, maximum variation of the force constant observed upon pairwise comparison all of the six crystallographic structures considered for human DHFR (, with the red line indicating the threshold value for selective mechanically sensitive residues.

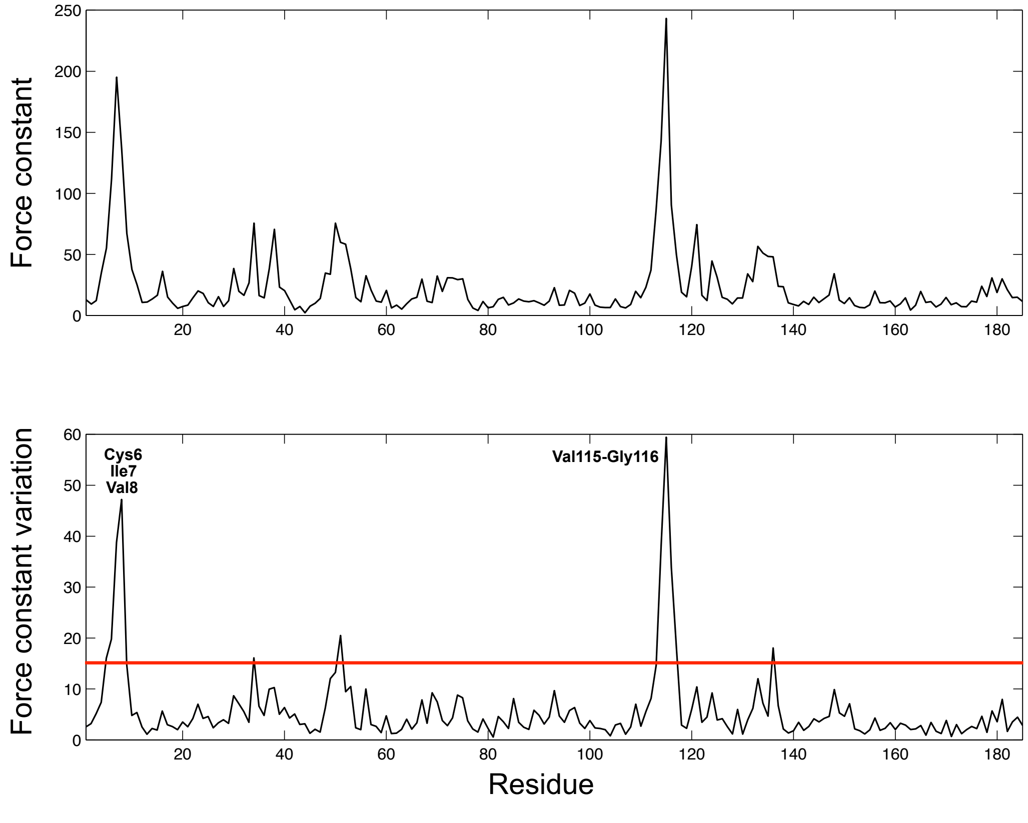

**Figure SI-3:** (a) Upper black line, rigidity profile for the crystallographic structure of human cyclophilin A (pdb 5noq). Lower black line, maximum variation of the force constant observed upon pairwise comparison all of the 10 crystallographic structures considered for human CPA. Lower red line, maximum variation of the force constant observed when considering only structures from 5noq, 5nos, 5nou and 5nov. (b) Distribution of the maximum force constant variation observed between two CPA structures as function of their Cα rmsd : Black points for all 10 crystallographic structures, red points for the structures from 5noq, 5nos, 5nou and 5nov only.

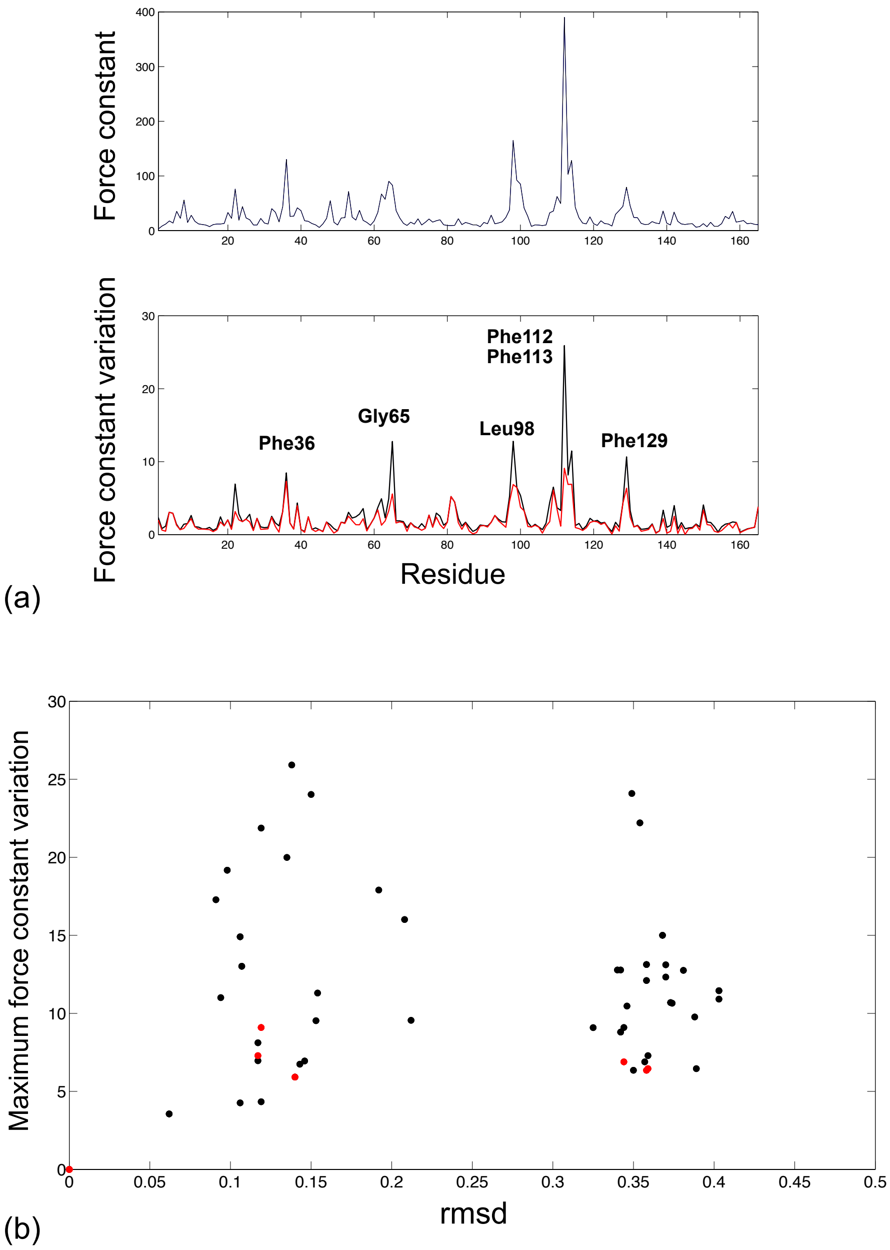

**Figure SI-4:** (a) Upper black line, rigidity profile of the first NMR representative model of DHFR from *B. anthracis* (pdb 2kgk). Lower black line, maximum variation of the force constant observed upon pairwise comparison all of the NMR models from the pdb entry. Lower red line, maximum variation of the force constant observed upon pairwise comparison of the representative models listed in Table 1 only . (b) Distribution of the maximum force constant variation observed between two DHFR structures as function of their Cα rmsd : Black points for all 15 models in the 2kgk pdb entry, red points for the 6 representative models listed in Table 1 only.

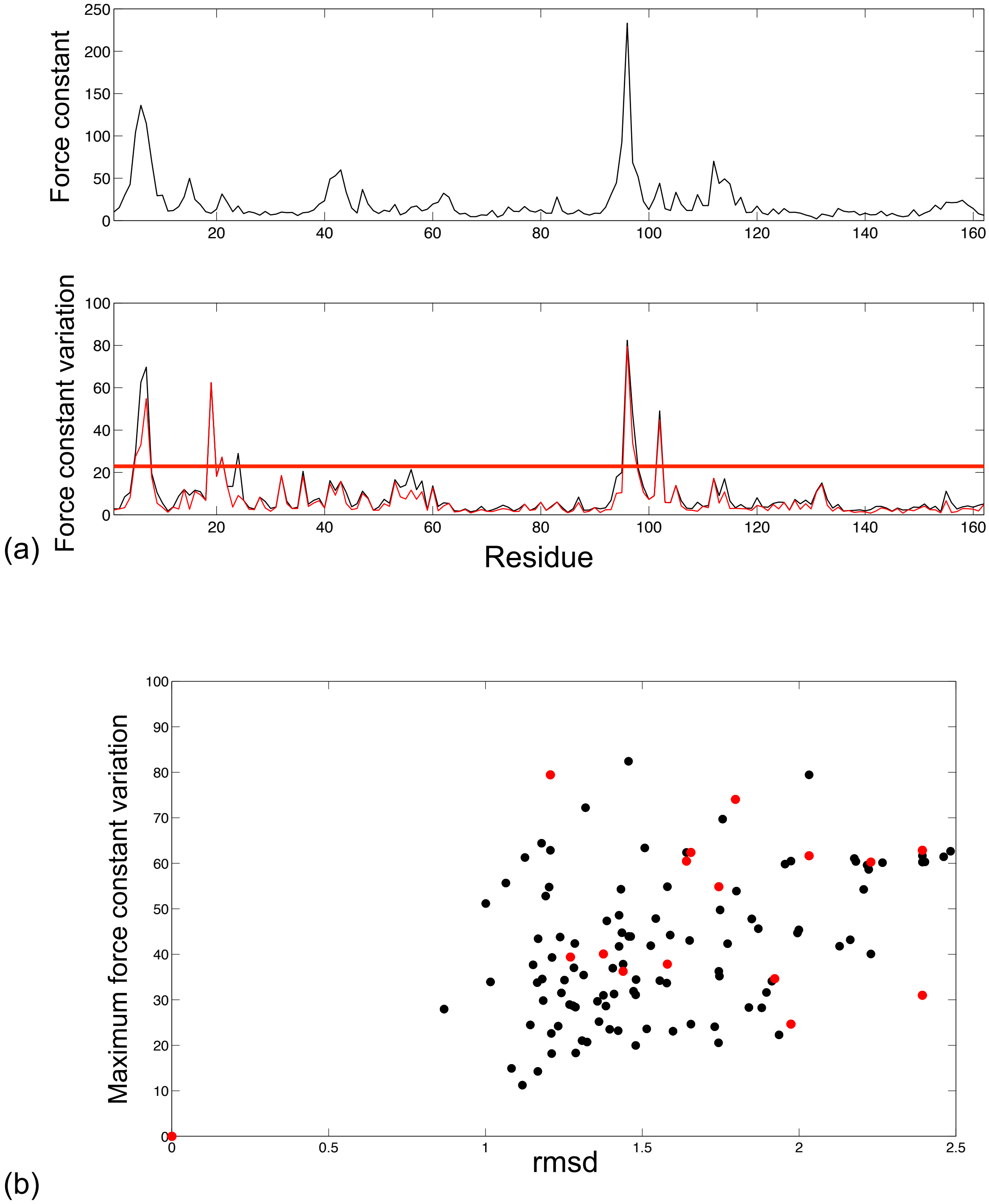
